# Microbiota-derived butyrate promotes dermal collagen production through epidermal-dermal crosstalk

**DOI:** 10.64898/2026.06.15.731906

**Authors:** Jamie Ting-Chun Pan, Sofía M. Murga-Garrido, Deborah Minzaghi, Ellen K. White, Sixia Huang, Aayushi Uberoi, Elizabeth A. Grice

## Abstract

The commensal microbiota actively shapes skin homeostasis by regulating immune responses and barrier function, yet interactions with the dermis and dermal fibroblasts remain poorly defined. Here, we show that the commensal microbiota regulates dermal collagen homeostasis through epidermal-mediated signaling. Transcriptomic analysis revealed enrichment of extracellular matrix (ECM)-associated gene programs in the dermis of conventionally raised (CR) mice compared with germ-free (GF) counterparts. CR skin exhibited increased collagen content, a thicker dermis, and thicker collagen fibers, indicative of a structurally robust dermal ECM. Similar microbiota-dependent collagen regulation was observed in additional tissues and during embryonic skin development, suggesting contributions from microbiota-derived signals beyond directly colonized sites. Mechanistically, short chain fatty acid butyrate robustly induced epidermal platelet-derived growth factor B (PDGFB) expression *in vitro* through a histone acetylation-dependent transcriptional program. Conditioned media from butyrate-treated human epidermal equivalents promoted collagen production in primary human dermal fibroblasts, an effect that was attenuated by inhibition of PDGF receptor signaling or histone acetyltransferase activity. Moreover, topical butyrate treatment of explants derived from aged human skin induced epidermal PDGFB and dermal collagen expression. Together, these findings identify a microbiota-epidermis-dermis signaling axis in which the commensal microbiota regulates dermal collagen through metabolite-enhanced epidermal signaling, highlighting a previously underappreciated role for the microbiota in maintaining dermal homeostasis.

## Introduction

The skin forms the first line of defense to the external environment, preventing the penetration of toxic and pathogenic substances, while protecting the body’s organs and preventing moisture loss. These critical functions rely on coordination across multiple cell types and layers. The epidermis, comprised of keratinocytes, forms the outermost layer of skin and the permeability barrier^1^. Beneath this, the dermal layer is composed of fibroblasts that deposit and remodel an extracellular matrix (ECM), thereby supporting tissue architecture and repair while dynamically regulating the cellular microenvironment^2^. In recent years, a role for the microbiota has emerged in regulating various facets of skin’s barrier function, including antimicrobial and tolerogenic immune responses, keratinocyte differentiation and epithelialization, and sebum production^3^.

Microbial metabolites and other products of microbial cells are emerging as chemical mediators that can regulate host cellular and molecular processes^4^. Amino acid metabolites, such as tryptophan metabolites produced by skin commensal bacteria, stimulate keratinocyte differentiation and attenuate atopic-like inflammation via ligand-receptor interaction with the aryl hydrocarbon receptor^5–7^. Short chain fatty acids (SCFA; eg. butyrate, propionate, acetate) are produced by gut microbiota-mediated fermentation of dietary fiber, whereas skin commensals such as *Cutibacterium acnes* and *Staphylococcus epidermidis* ferment sebum and glycerol to SCFAs under anaerobic conditions^8,9^. SCFAs act on host cells directly by binding G-protein-coupled fatty acid receptors, or through epigenomic effects acting as histone deacetylase (HDAC) inhibitors or enhancing histone acetylation^10^. Locally, SCFA can suppress skin inflammation via keratinocyte-directed^11,12^ and T regulatory cellular responses^13^, but in other contexts can be pro-inflammatory^14,15^. Recent evidence has highlighted how SCFA produced intestinally by the gut microbiota enter circulation and traffic to the skin to modulate keratinocyte metabolism and differentiation, and to limit allergen sensitization^16^. Thus, microbial metabolites produced locally by skin microbiota, or distally by the intestinal microbiota, can impact homeostatic skin barrier function and responses to inflammatory stimuli.

While microbial metabolites, such as SCFAs, have been shown to modulate keratinocyte and immune cell activity, their impact on dermal fibroblasts is less clear. Dermal fibroblasts synthesize, organize, and maintain a collagen-rich ECM that provides the skin its extraordinary tensile strength and a substrate for epidermal keratinocyte migration and stratification during development and wound healing^17^. Type I fibrillar collagen is highly abundant in the ECM, along with fibronectin, elastin, laminins, and other ‘matrisome’ proteins^18^. ECM synthesis, deposition, and remodeling is dynamic and regulated by growth factors including transforming growth factor-beta (TGFβ), platelet-derived growth factors (PDGFs), and epidermal growth factors (EGFs)^19^. Excessive collagen deposition causes fibrotic skin conditions, such as scleroderma, hypertrophic scarring, and keloids^20^. On the other hand, with aging, dermal fibroblasts progressively acquire a senescent phenotype characterized by reduced collagen production and increased expression of proteolytic and inflammatory factors^21^. Senescent fibroblasts, and altered collagen homeostasis, leads to dermal thinning and an altered ECM microenvironment that is linked to impaired wound healing, frailty, aging, and epithelial cancer development^22^. Thus, type I collagen homeostasis plays a critical role in the skin’s integrity and requires regulation of biosynthetic and degradative processes.

Here we investigated the role of the microbiota in regulating dermal fibroblast function and collagen homeostasis. Using gnotobiotic mouse models, we found that the microbiota regulates the composition and organization of the collagen fibers in the ECM. These effects were dependent on circulating microbial metabolites, and became apparent prenatally when maternal microbiota was disrupted during pregnancy. The SCFA microbial metabolite butyrate was found to be a potent inducer of PDGFB in human keratinocytes, which then stimulated human fibroblasts to produce collagen via paracrine signaling. In this context, we uncovered a mechanism where butyrate induces a metabolic-epigenetic program in keratinocytes to promote PDGFB expression via acetylation-regulated transcription. Finally, we show that topical delivery of butyrate to aged human skin can promote both epidermal PDGFB and dermal collagen production, suggesting a mechanism that can be therapeutically targeted to restore age-related ECM collagen homeostasis. Our findings reveal a mechanistic role for the microbiota in mediating a keratinocyte-fibroblast axis that regulates dermal collagen production and organization, and uncovers a potential therapeutic target with implications for fibrotic and senescence-related disorders of the skin.

## Results

### Commensal microbiota regulates the dermal matrisome

To investigate effects of the microbiota on dermal gene expression programs in intact skin, we performed bulk RNA sequencing (RNA-seq) on dermal compartments isolated from specific pathogen-free, conventionally raised (CR, n = 5) and germ-free (GF, n = 5) 12-week-old C57BL6/J mice (**Fig. 1A**). A third group of GF mice was topically colonized for 4 weeks with a defined human skin microbiota consortium, Flower’s Flora 50 (FF, n = 5), as previously described^6,23^ (**Fig. S1**). A total of 998 genes were differentially expressed between CR and GF dermis (**Fig. 1B**), compared to 247 between FF and GF (**Fig. S1A**). Among these, only 88 differentially expressed genes (DEGs) overlapped in the CR:GF and FF:GF analyses (**Fig. S1A**), suggesting that short-term skin colonization resulted in limited restoration of the GF dermal transcriptome. Among the 998 CR:GF DEGs, 519 were upregulated in CR while 479 were upregulated in GF dermis (**Fig. 1B**). We mapped these DEGs to a curated matrisome database of extracellular matrix (ECM) core proteins and regulatory factors, organized into six major categories^24^. Genes upregulated in CR showed striking enrichment in matrisome components, outnumbering GF DEGs across most categories (**Fig. 1C**). Notably, genes in the Collagens category showed the most contrast, and we therefore highlighted CR-enriched DEGs involved in collagen biosynthesis pathways (**Fig. 1D**). Although not statistically significant, collagen biosynthesis genes were partially restored in dermis from FF-colonized mice (**Fig. S1B**), suggesting that the extent of matrisome gene recovery may depend on the nature and duration of the microbial intervention. Together, these findings indicate that the commensal microbiota may modulate dermal matrisome composition.

**Figure 1.**
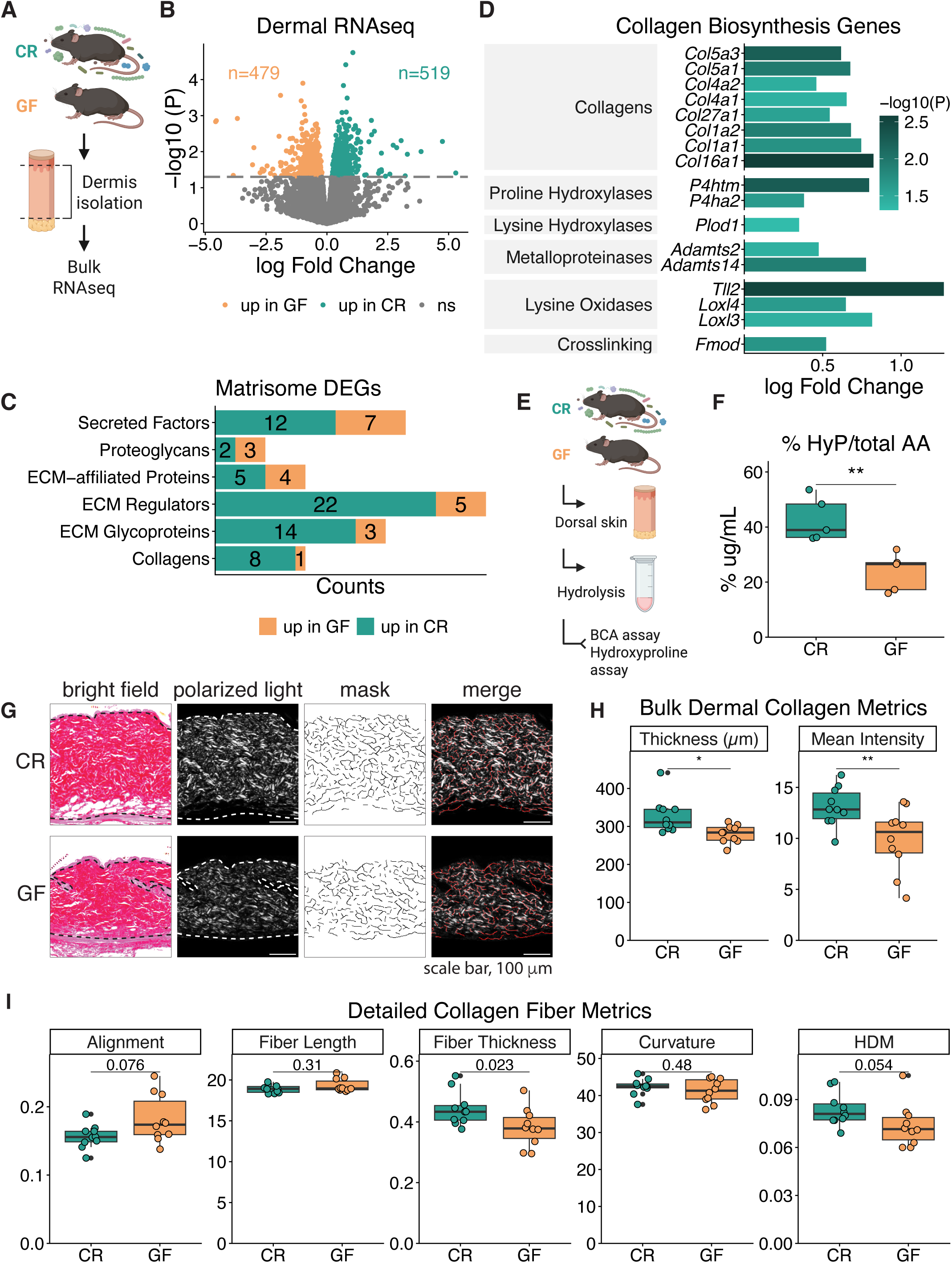
Commensal microbiota regulates dermal matrisome composition and collagen organization. (A) Schematic illustrating RNA-seq performed on dermal tissue isolated from tail skin of conventionally-raised (CR) and germ-free (GF) mice. N = 5 mice per group. (B) Volcano plot showing differentially expressed genes (DEGs) in CR versus GF dermis. The dashed line indicates a P-value threshold of 0.05. (C) Counts of matrisome genes in each category enriched among DEGs in CR and GF dermis. (D) CR-enriched DEGs associated with collagen biosynthesis pathways. Gray boxes denote general protein categories corresponding to the listed DEGs. (E) Workflow illustrating hydroxyproline (HyP) assay. HyP is quantified following alkaline hydrolysis and normalization to total amino acids (AA) measured by bicinchoninic acid (BCA) assay. (F) Hydroxyproline content in dorsal skin from CR and GF mice. N = 5 mice per group. **, P < 0.01 by Wilcoxon test. (G) Representative cropped picrosirius red (PSR) stained images of CR and GF dorsal skin shown as bright field, polarized light, TWOMBLI generated masks, and merged polarized light and mask images. Dashed lines denote the boundaries of dermal collagen signals. In polarized light images, signal is shown within manually defined dermal regions. N = 10 mice per group. Scale bar, 100 µm. (H) Dermal thickness and mean staining intensity quantified from polarized light images across the entire tissue section. *, P < 0.05; **, P < 0.01 by Wilcoxon test. (I) Selected collagen fiber metrics quantified from TWOMBLI generated masks across the entire tissue section. P-values were determined by Wilcoxon test.

To validate the transcriptomic findings at the tissue level, we quantified hydroxyproline (HyP) content in dorsal skin tissues isolated from CR and GF mice. HyP is a post-translationally modified amino acid enriched in fibrillar collagens that is assayed as a measure of total collagen content^25^. Consistent with the transcriptional data, HyP levels were significantly higher in CR skin compared to GF skin, after normalization to total amino acids (**Fig. 1E-F**). We next assessed collagen organization by histological staining of dorsal skin sections with picrosirius red (PSR). Under polarized light, PSR-stained thicker type I collagen fibers appear red/orange, whereas thinner type III collagen fibers appear green^26^. Given that collagen I predominates in uninjured, normal skin^27^ and that both *Col1a1* and *Col1a2* were transcriptionally upregulated in CR dermis (**Fig. 1D**), we extracted the red channel from polarized light images and converted to greyscale for analysis (**Fig. 1G**, polarized light). CR dermis was significantly thicker and higher in collagen staining intensity, compared to GF dermis (**Fig. 1H**). To further quantify collagen fiber organization, we applied TWOMBLI, an image analysis pipeline that extracts multiple metrics of matrix patterning^28^ (**Fig. 1G**, mask). While collagen fibers from CR and GF skin showed comparable fiber length and curvature, CR dermis exhibited significantly greater fiber thickness, along with a trend toward increased matrix density, measured as high-density matrix (HDM), and reduced fiber alignment, a feature consistent with a mechanically resilient matrix (**Fig. 1I**). Together, these features indicate that microbial colonization is associated with a thicker and more structurally robust dermal collagen matrix at baseline.

### Fibroblasts derived from germ-free skin are impaired in collagen production and migratory capacity

To investigate whether the collagen phenotype extends to the cellular level, we derived primary dermal fibroblasts from CR and GF mice (n = 5 per group). Cells were pooled within each condition to assess cell-associated collagen production by western blot and HyP assay, as well as collagen secreted into the culture medium using an in-solution PSR assay^29^. All three assays revealed higher collagen levels in CR-derived fibroblasts compared to GF fibroblasts (**Fig. S2A-C**), consistent with the tissue-level results. In parallel, fibroblast migratory capacity was assessed using dermal fibroblasts derived from individual mice, revealing greater migration of CR-derived fibroblasts in a scratch closure assay (**Fig. S2D-E**). To assess whether ECM derived from GF fibroblasts is functionally impaired, we next performed matrix-swapping assays in which recipient fibroblasts were seeded onto ECM deposited by donor fibroblasts following decellularization (**Fig. S2F**). While intrinsic cellular differences remained prominent, ECM derived from CR fibroblasts significantly improved migration of GF recipient fibroblasts, whereas GF-derived ECM impaired migration of CR recipient fibroblasts (**Fig. S2G**). Together, these results indicate that the dermal collagen differences observed *in vivo* are retained in isolated dermal fibroblasts.

### Microbial metabolites regulate dermal collagen indirectly

To begin dissecting the nature of the microbial exposure that mediates dermal collagen production, we first analyzed the tissue specificity of the phenotype. We therefore assessed collagen content in intestine, a microbiota-associated tissue, and tail bone, a collagen-rich but non-microbiota-associated tissue. Collagen levels were significantly increased in CR intestine compared to GF intestine, whereas tail bone collagen exhibited a consistent upward trend in CR mice, albeit not reaching statistical significance (**Fig. 2A**). These data suggest that microbiota-dependent regulation of collagen may extend beyond directly colonized tissues, potentially via circulating microbiota-derived signals.

**Figure 2.**
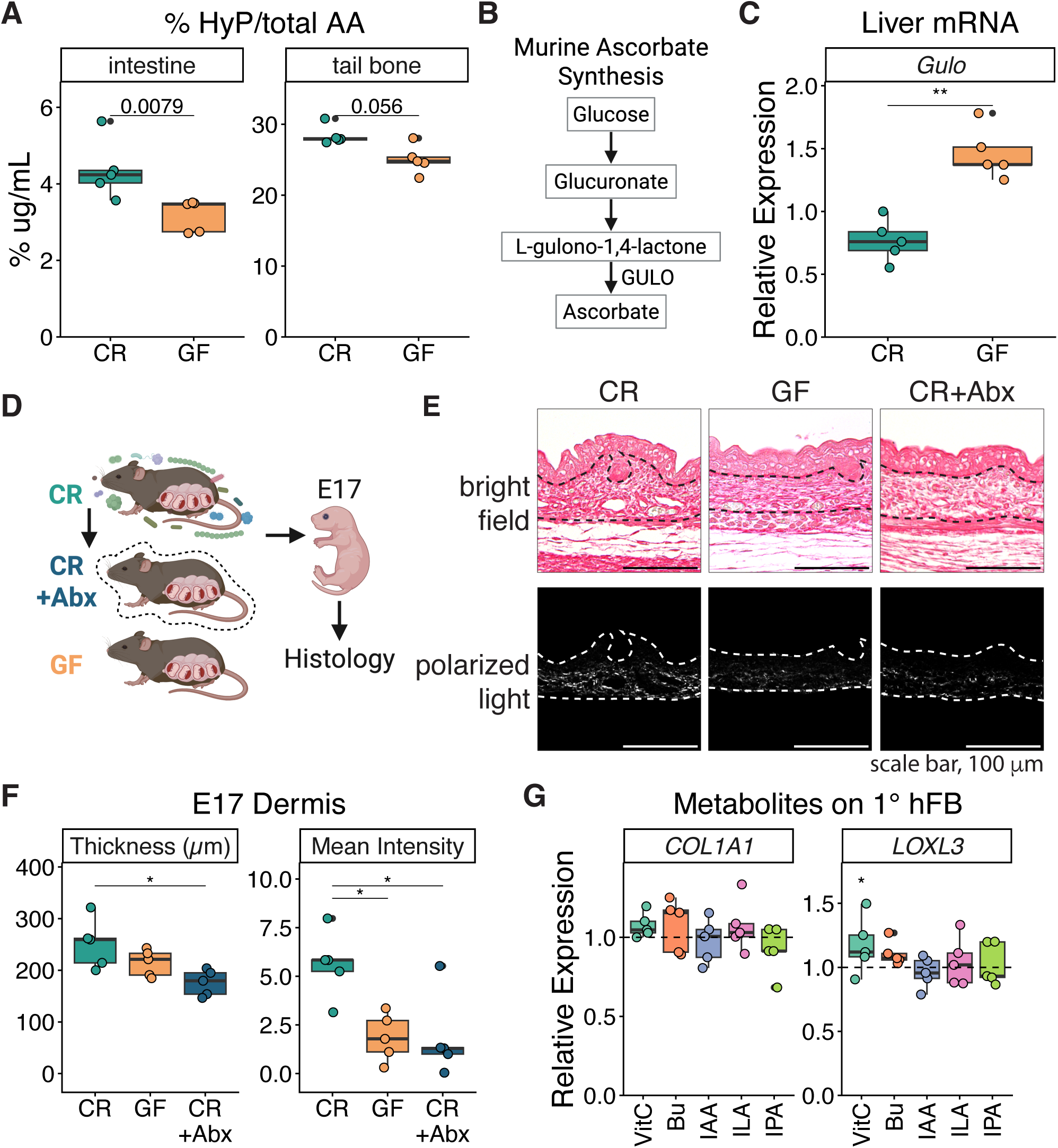
Microbial metabolites regulate dermal collagen indirectly. (A) HyP content of intestine and tail bone tissues isolated from CR and GF mice normalized to total AA. P-values were determined by Wilcoxon test. N = 5 mice per group. (B) Simplified murine ascorbate synthesis pathway. (C) Hepatic *Gulo* mRNA levels in CR and GF mice measured by qPCR and normalized to *Rplp2*. **, P < 0.01 by Wilcoxon test. N = 5 mice per group. (D) Embryonic skin was collected from E17 embryos harvested from CR, GF, and CR dams treated with systemic antibiotics (Abx). N = 5 dams per group; one embryo per dam. (E) Representative cropped picrosirius red (PSR) stained images of embryonic dorsal skin shown as bright field and polarized light images. Dashed lines denote boundaries of dermal collagen signals. In polarized light images, signal is shown within defined dermal regions. Scale bar, 100 µm. (F) Dermal thickness and mean staining intensity quantified from two fields per embryo from polarized light images. *, P < 0.05 by Wilcoxon test. (G) *COL1A1* and *LOXL3* mRNA levels normalized to *RPLP0* from primary human dermal fibroblasts (1° hFB) treated for 24 hours with vehicle control or selected microbial metabolites, including 200 µM ascorbic acid (VitC), 2 mM butyrate (Bu), 250 µM indole derivatives: indole-3-acetic acid (IAA), indole-3-lactic acid (ILA), and indole-3-propionic acid (IPA). Dashed lines denote vehicle control levels normalized to 1 for each donor. N = 5 donors. *, P < 0.05 by linear mixed effects model with donor as a random effect.

To exclude the possibility that GF mice might be deficient in endogenous ascorbic acid, a critical cofactor for collagen biosynthesis, we next examined hepatic expression of *Gulo*, which encodes gulonolactone oxidase, the rate-limiting enzyme for murine ascorbic acid synthesis^30^ (**Fig. 2B**). Unlike humans, mice retain the capacity for endogenous ascorbic acid production^30^. *Gulo* expression was significantly elevated in the livers of GF mice compared to CR mice (**Fig. 2C**), indicating that diminished collagen levels in GF mice are unlikely to be attributable to reduced endogenous ascorbic acid availability.

To assess whether circulating microbial metabolites influence dermal collagen independently of direct microbial colonization, and to establish timing of the exposure, we analyzed embryonic skin. The in utero environment is devoid of live microbes but remains exposed to maternal microbial metabolites via the placental interface^11–14^. Embryos were collected at embryonic day 17 (E17) from CR dams, GF dams, and CR dams treated with antibiotics during embryonic dermal morphogenesis (n = 5 dams per group; one embryo per dam; **Fig. 2D**). Embryonic skin from CR dams exhibited increased dermal thickness and collagen staining intensity compared to GF embryos (**Fig. 2E-F**), consistent with the phenotype observed in adult mice (**Fig. 1G-H**). Embryos from antibiotic-treated CR dams phenocopied GF embryos, displaying reduced dermal collagen thickness and intensity (**Fig. 2E-F**). Together, these observations support a role for circulating microbiota-derived metabolites in shaping dermal collagen production and organization.

We next tested whether microbial metabolites directly modulate dermal fibroblasts by treating primary human dermal fibroblasts (1° hFB) isolated from neonatal foreskin with a panel of selected microbial metabolites, including ascorbic acid (VitC), butyrate (Bu), indole-3-acetic acid (IAA), indole-3-lactic acid (ILA), and indole-3-propionic acid (IPA). These metabolites are microbially derived and/or have previously been reported to regulate collagen expression or influence skin biology^6,16,35–37^. VitC, a known cofactor for collagen biosynthesis that can also be microbially produced^38^, was included as a reference condition. With the exception of VitC, none of the tested metabolites induced significant or consistent changes in collagen expression, as assessed by the mRNA levels of *COL1A1* and *LOXL3*, a key enzyme involved in collagen crosslinking^39^ (**Fig. 2G**). These results suggest that metabolite-mediated modulation of dermal collagen occurs indirectly.

### Commensal microbiota facilitates epidermal-dermal crosstalk

Because the epidermis is the primary interface for local microbiota-derived factors and collagen expression is frequently stimulated through paracrine signaling^2,3,40–42^, we hypothesized that microbiota-dependent regulation of dermal collagen is mediated indirectly through epidermal-dermal crosstalk. To identify potential epidermal-dermal interactions facilitated by the microbiota, we merged dermal DEGs (**Fig. 1B**) with DEGs extracted from our previously published epidermal RNA-seq dataset^23^ (n = 4 per group; **Fig. 3A**) to generate a combined DEG list for CR and GF skin. This combined DEG list was then used to query a ligand-receptor database from OmniPath^43^ (**Fig. 3B**). Despite comparable numbers of DEGs, CR skin exhibited markedly more dynamic inter- and intra-tissue signaling than GF skin (**Fig. 3C**). Detailed ligand-receptor interaction networks for CR and GF skin (**Fig. 3D-E**, respectively) illustrate the drastic reduction of predicted epidermal-dermal interactions in GF skin. Within the CR network, *TGFB1* and *PDGFB* emerged as top candidates based on their well-documented roles in fibroblast biology and collagen production^2^ (**Fig. 3D**). Protein levels of both growth factors were significantly increased in CR compared to GF epidermis, as measured by ELISA (**Fig. 3F-G**). Collectively, these data support a model in which commensal microbiota enhances inter- and intra-tissue signaling in the skin, with epidermal-derived growth factors as candidate mediators of dermal collagen regulation.

**Figure 3.**
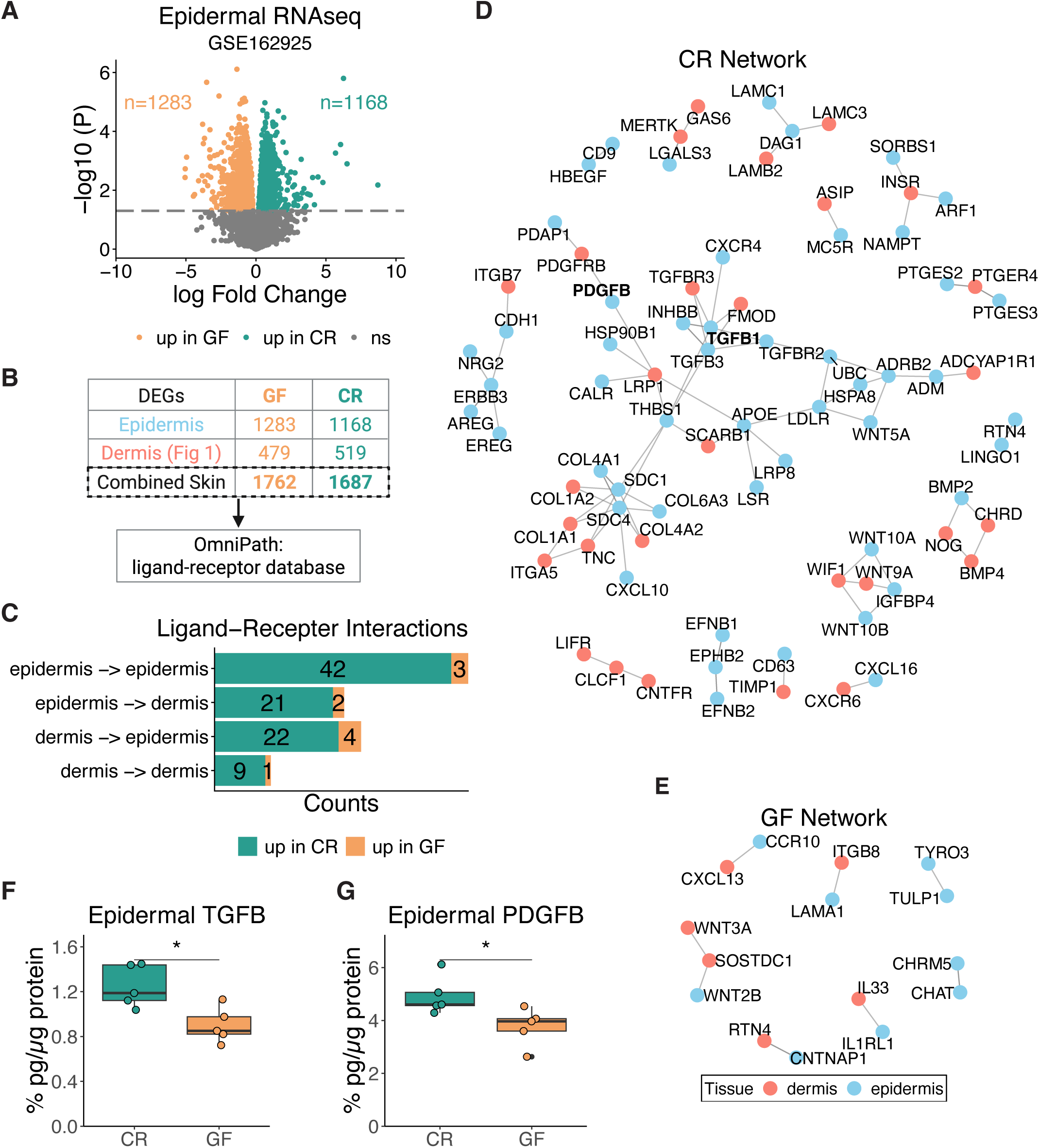
Commensal microbiota facilitates epidermal-dermal crosstalk. (A) Volcano plot showing DEGs in CR versus GF epidermis. Original data were obtained from GSE162925. N = 4 mice per group. The dashed line indicates a P-value threshold of 0.05. (B) Workflow illustrating integration of epidermal and dermal RNA-seq datasets to identify combined skin DEGs and map ligand-receptor interactions using OmniPath. (C) Counts of ligand-receptor pairs enriched among combined skin DEGs in CR and GF conditions, stratified by interaction direction between epidermis and dermis. (D-E) Ligand-receptor interaction networks within (D) CR and (E) GF skin derived from combined epidermal and dermal DEGs. (F-G) (F) Latent TGFβ and (G) PDGFB protein levels in CR and GF epidermis quantified by ELISA and normalized to total protein. N = 5 mice per group. *, P < 0.05 by Wilcoxon test.

### Butyrate promotes epidermal PDGFB signaling to induce dermal collagen expression

To test whether microbial metabolites modulate epidermal production of candidate paracrine factors, we treated primary human keratinocytes (1° hKC) with a panel of microbial-derived metabolites as before. In undifferentiated monolayer culture, ILA modestly increased *TGFB1* expression, whereas Bu significantly induced *PDGFB* expression (**Fig. 4A**). We next evaluated metabolite responses in a three-dimensional setting, where human epidermal equivalents (HEE) were generated by culturing 1° hKC on transwells followed by air-liquid interface differentiation, with metabolites supplied in the culture media (**Fig. 4B**). In contrast to the monolayer model, ILA did not induce *TGFB1* expression in the HEE. Instead, Bu robustly increased *PDGFB* expression by up to fivefold relative to vehicle control, accompanied by increased expression of the differentiation marker involucrin (*IVL*) and reduced *TGFB1* expression (**Fig. 4C**). Histological analysis confirmed stratified epidermal architecture and revealed morphology consistent with accelerated differentiation in Bu-treated HEEs, including reduced basal-like cell prominence and thickened stratum corneum (**Fig. 4D**). IPA modestly induced *PDGFB* expression but did not increase *IVL*, in contrast to Bu, which induced both *PDGFB* and differentiation markers (**Fig. 4D**).

**Figure 4.**
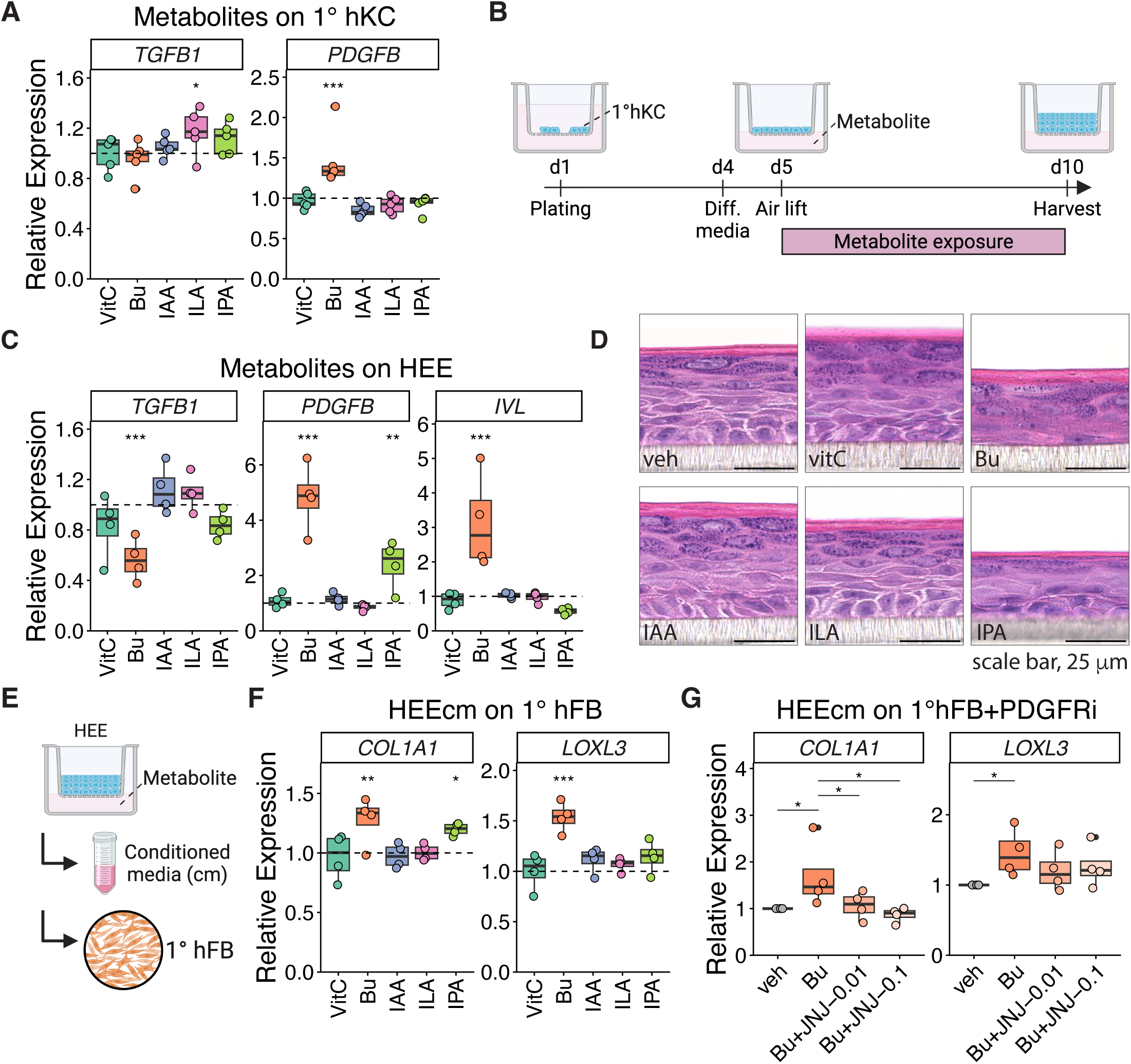
Butyrate promotes epidermal PDGFB signaling to induce dermal collagen expression. (A) *TGFB1* and *PDGFB* mRNA levels normalized to *RPLP0* from primary human keratinocytes (1° hKC) treated with vehicle control or indicated microbial metabolites at the same concentrations as in Fig. 2D for 4 hours. Dashed lines denote vehicle control levels normalized to 1 for each donor. N = 5 donors. *, P < 0.05; ***, P < 0.001 by linear mixed effects model with donor as a random effect. (B) Schematic illustrating human epidermal equivalent (HEE) cultures. Primary hKC were plated on transwells, switched to differentiation media (Diff. media), and air lifted for differentiation prior to treatment with vehicle control or microbial metabolites. (C) *TGFB1*, *PDGFB*, and *IVL* mRNA levels normalized to *RPLP0* from HEEs treated with vehicle control or indicated microbial metabolites at 100 µM VitC, 1 mM Bu, and 125 µM indole derivatives. Dashed lines denote vehicle control levels normalized to 1 for each donor. N = 4 donors. *, P < 0.05; ***, P < 0.001 by linear mixed effects model with donor as a random effect. (D) Hematoxylin and eosin stained HEEs treated as indicated. Scale bar, 25 µm. (E) Workflow illustrating treatment of 1° hFB with conditioned media (cm) collected from metabolite treated HEEs. (F) COL1A1 and *LOXL3* mRNA levels normalized to *RPLP0* from 1° hFB treated with HEE cm as indicated for 24 hours. Dashed lines denote vehicle control levels normalized to 1 for each donor. N = 4 HEE donors. **, P < 0.01; ***, P < 0.001 by linear mixed effects model with donor as a random effect. (G) *COL1A1* and *LOXL3* mRNA levels normalized to *RPLP0* from 1° hFB treated with vehicle or 1 mM Bu treated HEE cm in the presence of a PDGF receptor inhibitor, JNJ-10198409 (PDGFRi, JNJ, Bu+JNJ) at 0.01 or 0.1 µM for 24 hours, expressed relative to the corresponding vehicle control (veh) for each donor. N = 4 HEE donors. *, P < 0.05; **, P < 0.01 by linear mixed effects model with donor as a random effect.

To assess whether Bu-induced keratinocyte-derived PDGFB promotes dermal collagen expression, we treated 1° hFB with conditioned media (20%) collected from metabolite-treated HEEs (**Fig. 4E**). Conditioned media from Bu-treated HEEs significantly increased expression of both *COL1A1* and *LOXL3* in 1° hFB (**Fig. 4F**), consistent with elevated epidermal *PDGFB* expression and PDGFB abundance following Bu treatment (**Fig. 4C, Fig. S3B**). Conditioned media from IPA-treated HEEs modestly increased *COL1A1* expression alone (**Fig. 4F**), consistent with its weaker induction of epidermal *PDGFB* expression (**Fig. 4C**). To test whether the Bu-induced fibroblast response was mediated through PDGFB signaling, we blocked PDGF receptor (PDGFR) activity during fibroblast stimulation. PDGFR inhibition by the selective inhibitor JNJ-10198409 (JNJ) markedly attenuated Bu-mediated induction of *COL1A1* and *LOXL3* (**Fig. 4G**). Together, these results indicate that Bu-enhanced keratinocyte-fibroblast crosstalk promotes collagen expression through PDGFB signaling.

Bu-induced *PDGFB* expression in the HEE was dose dependent and positively correlated with epidermal differentiation (**Fig. S3A-B**). Expression of the basal keratinocyte marker keratin 14 (*KRT14*) was reduced, while expression of differentiation markers keratin 10 (*KRT10*), *IVL*, loricrin (*LOR*), and filaggrin (*FLG*) were increased (**Fig. S3A**). These findings were corroborated by histological analysis of HEE constructs, where increasing Bu concentration resulted in visibly increased differentiation (**Fig. S3B**). PDGFB protein expression was further validated by immunostaining, with intracellular PDGFB signal in keratinocytes and additional signal associated with the transwell membrane, potentially reflecting accumulation of secreted PDGFB at the epithelial interface (**Fig. S3B**).

The accelerated keratinocyte differentiation induced by Bu was consistent with previous reports^16^. Therefore, we next determined whether Bu-induced *PDGFB* expression was dependent on keratinocyte differentiation state by using a calcium-induced monolayer differentiation model. As expected, high calcium alone increased expression of both the early and late differentiation markers *KRT10* and *IVL*, with the addition of Bu further enhancing expression of the late marker *IVL* (**Fig. S3C**). In contrast, *PDGFB* expression was robustly induced by Bu regardless of calcium concentration (**Fig. S3C**), indicating that PDGFB induction is Bu-specific and independent of keratinocyte differentiation state.

To determine whether *PDGFB* induction was specific to Bu or shared among short-chain fatty acids (SCFAs), we evaluated the effects of acetate and propionate on 1° hKC. Propionate modestly induced *PDGFB* expression (**Fig. S3E**), whereas acetate showed minimal or no induction compared to 1 mM Bu, even at higher concentrations (**Fig. S3D**). Propionate-induced *PDGFB* expression remained weaker than that observed with Bu, indicating that SCFAs differ in their capacity to promote epidermal *PDGFB* signaling. These findings highlight Bu as the most potent SCFA driving epidermal *PDGFB* expression.

### Butyrate induces epidermal PDGFB via a metabolic-epigenetic mechanism

We next sought to elucidate the mechanism by which Bu induces *PDGFB* expression. Because Bu is a well-characterized histone deacetylase (HDAC) inhibitor^44^, we tested whether HDAC inhibition alone was sufficient to induce *PDGFB* in the HEE model. Treatment with suberoylanilide hydroxamic acid (SAHA, vorinostat), an HDAC inhibitory drug, only modestly induced *PDGFB* at the highest concentration, and did not reach levels comparable to Bu, despite robust, dose-dependent induction of the differentiation marker *IVL* (**Fig. 5A-B**). As a positive control for HDAC-dependent Bu responses^45^, *AQP3* expression was induced by both Bu and SAHA, confirming effective HDAC inhibition in this system (**Fig. S4A**). Consistent with this, SAHA-treated HEEs exhibited morphology similar to Bu-treated HEEs, with features of advanced differentiation (**Fig. S4B**), further uncoupling PDGFB induction from epidermal differentiation. In addition, Bu can act as a ligand for select G protein-coupled receptors (GPCRs)^44^. However, treatment of HEEs with inhibitors of Gi/o (pertussis toxin, PTX) or Gq/11 (YM-254890, YM) signaling did not attenuate Bu-induced *PDGFB* expression (**Fig. S4C-E**). We also examined whether MAPK signaling contributes to Bu-induced *PDGFB* expression, as MAPK pathways have been implicated in Bu responses and PDGFB regulation, separately^46–48^; inhibition of MAPK pathways had no effect on *PDGFB* induction (**Fig. S4F**). Together, these results indicate that Bu-induced *PDGFB* expression occurs independently of commonly implicated butyrate signaling pathways, including HDAC inhibition, GPCR-mediated signaling, and MAPK pathways.

**Figure 5.**
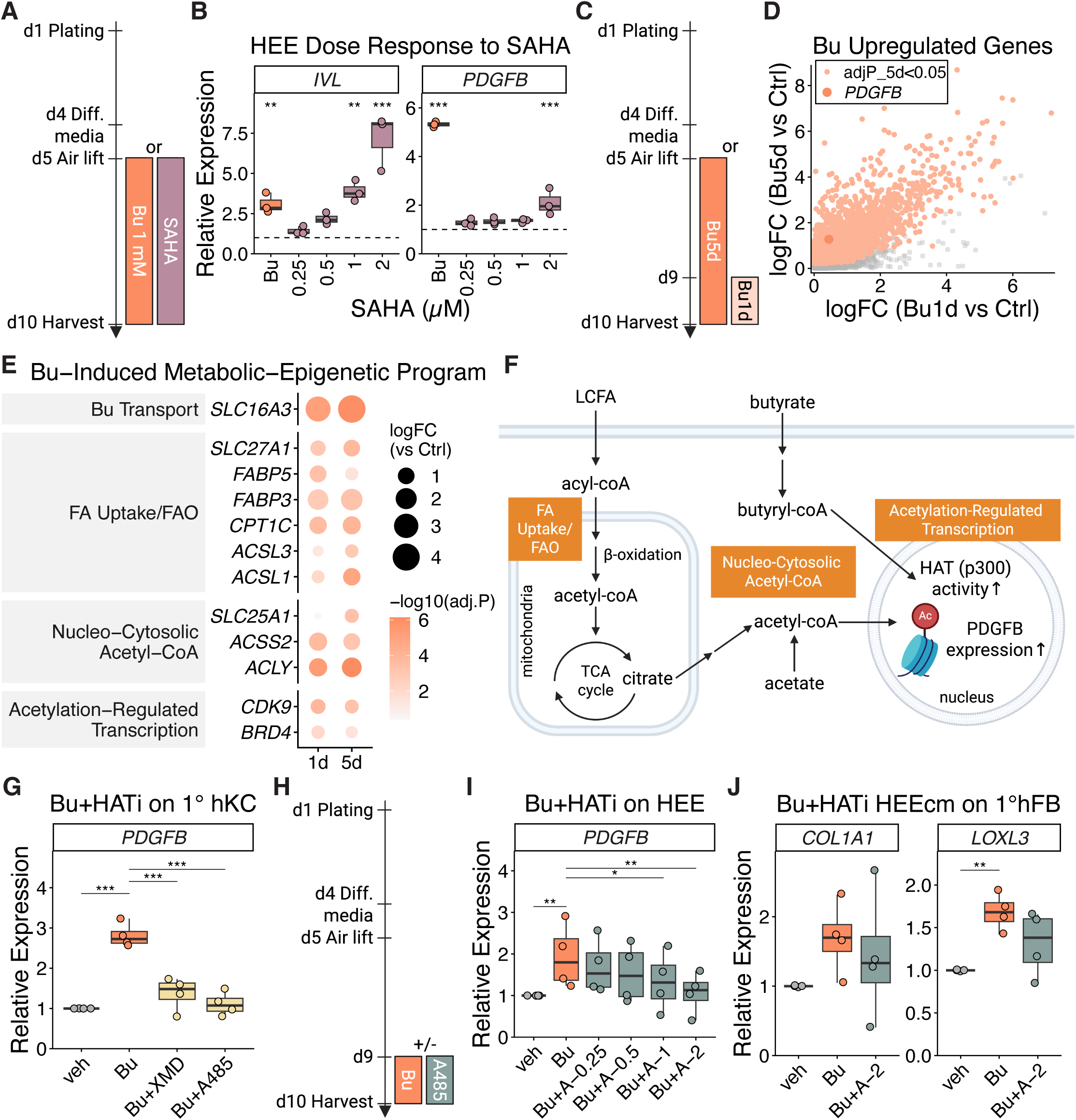
Butyrate induces epidermal PDGFB through a metabolic-epigenetic acetyl-CoA-HAT axis. (A) Workflow illustrating treatment of HEEs with either 1 mM Bu or suberoylanilide hydroxamic acid (SAHA). (B) *IVL* and *PDGFB* mRNA levels normalized to *RPLP0* from HEEs treated with vehicle, Bu or indicated concentrations of SAHA. Dashed lines denote vehicle control levels normalized to 1 for each donor. N = 3 donors. **, P < 0.01; ***, P < 0.001 by linear mixed effects model with donor as a random effect. (C) Workflow illustrating treatment of HEEs with either 1 mM Bu starting at air lift (Bu5d) or one day prior to harvest (Bu1d). (D) Scatter plot showing DEGs with positive log fold change (FC) in both Bu5d versus control (y-axis) and Bu1d versus control (x-axis). (E) DEGs highlighted in (D) associated with indicated functional categories, shown in gray boxes. (F) Proposed pathways of Bu-induced *PDGFB* expression. (G) *PDGFB* mRNA levels normalized to *RPLP0* from 1° hKC treated for 24 hours with vehicle, 1 mM Bu, Bu plus 2 µM BRD4 inhibitor (XMD8-92, Bu+XMD), or Bu plus 1 µM p300 inhibitor (A485, Bu+A485), expressed relative to the corresponding vehicle control for each donor. N = 4 donors. ***, P < 0.001 by linear mixed effects model with donor as a random effect. (H) Workflow illustrating treatment of HEEs with 1 mM Bu in the presence or absence of A485 one day prior to harvest. (I) *PDGFB* mRNA levels normalized to *RPLP0* from HEEs treated for 24 hours with vehicle or 1 mM Bu in the presence of indicated concentrations of A485, expressed relative to the corresponding vehicle control (veh) for each donor. N = 4 donors. *, P < 0.05; **, P < 0.01 by linear mixed effects model with donor as a random effect. (J) *COL1A1 and LOXL3* mRNA levels normalized to *RPLP0* from 1° hFB treated for 24 hours with vehicle or conditioned media collected from HEEs treated with 1 mM Bu in the presence or absence of 2 µM A485, expressed relative to the corresponding vehicle control (veh) for each donor. N = 4 HEE donors. **, P < 0.01 by linear mixed effects model with donor as a random effect.

To obtain a holistic view of transcriptional changes induced by Bu in the epidermis, we performed RNA-seq on HEEs treated with Bu either for five days following airlift (Bu5d) or for one day prior to harvest (Bu1d), alongside vehicle-treated controls (n = 4 per group; **Fig. 5C**). *PDGFB* was positively regulated at both Bu1d and Bu5d, with stronger induction at Bu5d (**Fig. 5D**). We reasoned that genes mediating Bu-induced *PDGFB* expression would exhibit induction kinetics similar to *PDGFB*. Candidate mediators were therefore defined as genes with positive log fold change in both Bu1d and Bu5d relative to control, and an adjusted P value less than 0.05 for Bu5d. Genes meeting these criteria are highlighted in **Fig. 5D**. Pathways involved in Bu transport as well as fatty acid (FA) uptake and beta-oxidation (FAO) were enriched (**Fig. 5E**). This pattern is consistent with prior reports showing that Bu enhances FA uptake and supports FAO, which generates acetyl-CoA to fuel the tricarboxylic acid (TCA) cycle^16,49^ (**Fig. 5F**). In addition, genes involved in increasing the nucleo-cytosolic acetyl-CoA pool were enriched, including *SLC25A1*, encoding the mitochondrial citrate carrier (CIC) that exports citrate from the TCA cycle to the cytosol^50^, *ACLY*, the ATP-citrate lyase that converts cytosolic citrate into acetyl-CoA^51–53^, and *ACSS2*, acetyl-CoA synthetase 2, which converts cytosolic and nuclear acetate and potentially Bu into acetyl-CoA^49,51,54,55^ (**Fig. 5E-F**). The expanded nucleo-cytosolic acetyl-CoA pool provides substrate for histone acetyltransferase (HAT) activity^51–53,55^. Consistent with this, Bu has been reported to enhance p300 HAT activity through auto-acylation^56^, and components of the p300-dependent transcriptional machinery, including *BRD4* and *CDKS*^57,58^, were also enriched following Bu treatment (**Fig. 5E**). Collectively, these data outline a Bu-induced metabolic-epigenetic program that may drive *PDGFB* expression.

To functionally validate the metabolic-epigenetic model suggested by the transcriptomic analysis, we next tested whether perturbation of key nodes within this program affects Bu-induced *PDGFB* expression. Primary hKCs were co-treated with Bu and inhibitors targeting the p300-dependent transcriptional machinery. Inhibition of BRD4 with XMD8-92 (XMD) or p300 with A485 significantly attenuated Bu-induced *PDGFB* expression (**Fig. 5G**). Similarly, p300 inhibition with A485 dose-dependently reduced Bu-induced *PDGFB* expression in HEEs (**Fig. 5H-I**). Histological analysis showed that A485-treated HEEs retained overall stratified epidermal architecture despite reduced *PDGFB* expression (**Fig. S4G**). In line with reduced epidermal *PDGFB* expression, conditioned media from A485-treated HEEs partially attenuated fibroblast responses elicited by conditioned media from Bu-treated HEEs, with a more pronounced effect observed for *LOXL3* expression (**Fig. 5J**). Together, these findings support a role for acetylation-dependent transcription in mediating Bu-induced *PDGFB* expression and downstream keratinocyte-fibroblast signaling.

### Topical butyrate enhances epidermal PDGFB and dermal collagen responses in explants derived from aged human skin

To determine whether the Bu-induced epidermal-dermal signaling axis is preserved in human skin, we utilized explants derived from aged human skin cultured at the air-liquid interface with topical Bu treatment in the presence or absence of the p300 inhibitor, A485 (**Fig. 6A**). Following treatment, epidermis and dermis were separated for compartment-specific qPCR analysis. Topical Bu significantly increased *PDGFB* expression in the epidermis and trended toward increased *COL1A1* expression in the dermis, whereas co-treatment with A485 attenuated both responses (**Fig. 6B-C**). Consistent with these findings, immunostaining revealed increased epidermal PDGFB signal and enhanced dermal procollagen type I (proCOL1) staining following Bu treatment, both of which were reduced by co-treatment with A485 (**Fig. 6D**). Together, these findings indicate that topical Bu promotes epidermal *PDGFB* signaling and dermal collagen responses in aged human skin and are consistent with a role for p300-dependent transcription in this process.

**Figure 6.**
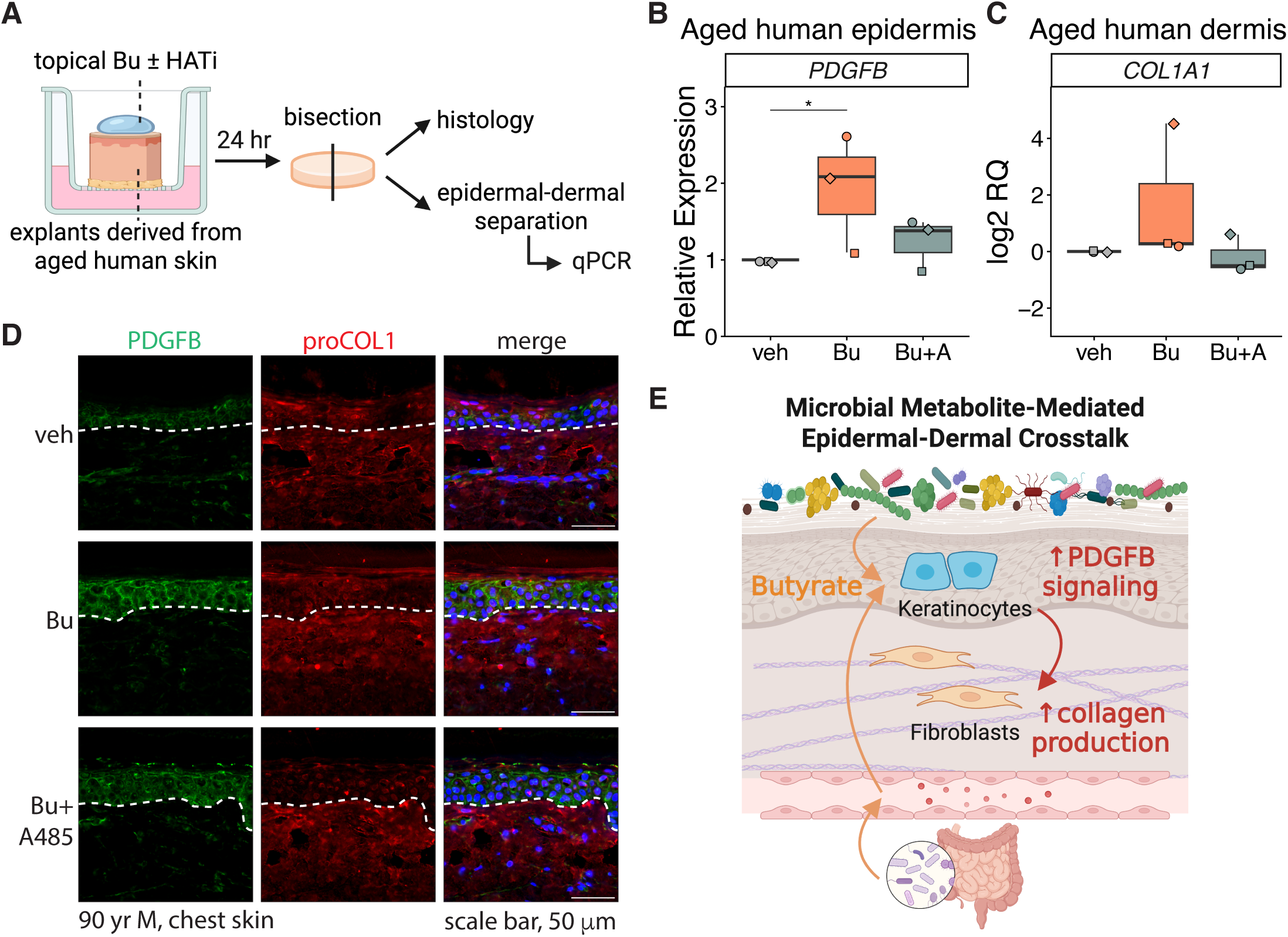
Topical butyrate enhances epidermal PDGFB and dermal collagen responses in explants derived from aged human skin. (A) Schematic illustrating processing of explants derived from aged human skin. Explants generated from aged human skin obtained during Mohs surgery were cultured at the air-liquid interface and topically treated with 2 mM Bu in the presence or absence of 5 µM A485 for 24 hours. Explants were then processed for histology or epidermal-dermal separation followed by qPCR analysis. (B-C) *PDGFB* and *COL1A1* mRNA levels normalized to *RPLP0* from the epidermal (B) and dermal (C) compartments of human skin explants treated with vehicle (veh), Bu, or Bu plus A485 (Bu+A). Data are expressed relative to the corresponding vehicle control for each donor. N = 3 donors. *, P < 0.05 by linear mixed effects model with donor as a random effect. (D) Representative immunostaining of PDGFB and procollagen type I (proCOL1) in explants derived from aged human skin treated as indicated. Dashed lines denote the epidermal-dermal junction. Nuclei were counterstained with DAPI. Scale bar, 50 µm. (E) Proposed model illustrating microbiota metabolite mediated epidermal-dermal crosstalk. Microbiota-derived butyrate promotes epidermal PDGFB signaling to enhance dermal collagen production.

## Discussion

The extent to which the microbiota contributes to tissue architecture beyond barrier surfaces remains poorly defined. Here, we uncover a previously unappreciated role for the microbiota in regulating dermal extracellular matrix (ECM) homeostasis through a keratinocyte-fibroblast signaling axis. Using gnotobiotic mouse models, we show that the commensal microbiota is required for establishing normal collagen composition and higher-order ECM organization **(Fig. 1)**. Notably, these effects are mediated by circulating microbial metabolites and are established early in development, as maternal microbiota disruption during pregnancy was sufficient to alter dermal collagen architecture in offspring **(Fig. 2)**. Mechanistically, we found that the short chain fatty acid butyrate (SCFA Bu) acts on keratinocytes to induce platelet-derived growth factor B (PDGFB) expression, which in turn promotes collagen production in dermal fibroblasts via paracrine signaling **(Fig. 4)**. This process is driven by a metabolic-epigenetic program in keratinocytes, whereby Bu enhances acetylation dependent transcriptional activation of PDGFB **(Fig. 5)**. Extending these findings to human tissue, we show that topical delivery of Bu to aged skin increases epidermal PDGFB and dermal collagen, suggesting that microbiota-derived metabolites can help restore age-associated dermal ECM decline **(Fig. 6)**. Together, we establish a mechanistic framework linking microbial metabolites to dermal structure and integrity, and identify a pathway that can be therapeutically targeted, with implications for dermal decline associated with aging and senescence.

Among candidate mediators of epidermal-dermal crosstalk, PDGFB emerged as a key epidermal-derived signal linking microbial metabolites to dermal collagen regulation (**Fig. 4**). PDGFs are well-established regulators of fibroblast activation and collagen production^2,59–61^, and keratinocytes have been reported to be a major source of PDGF in the skin while lacking expression for the receptor, PDGFR, themselves^62^. It is therefore striking that SCFAs induced epidermal PDGFB expression, seemingly to communicate with neighboring cell types. While our study focused on keratinocyte-fibroblast communication, PDGFB may also influence other dermal populations, including endothelial cells and immune cells, potentially affecting angiogenesis, vascular remodeling, and immune modulation^63–65^. These additional cell types may further contribute to extracellular matrix remodeling and dermal homeostasis. PDGFB can form both homodimers and heterodimers with PDGFA, and these distinct dimers may differentially regulate collagen production^66,67^. Notably, PDGFA was not upregulated in our transcriptomic datasets, suggesting that microbiota-dependent signaling may preferentially enhance PDGF-BB-driven pathways. Future studies incorporating more complex *in vivo* systems or single-cell approaches will help delineate these broader cellular contributions.

Our data further suggest that Bu induces PDGFB expression through a metabolic-epigenetic mechanism (**Fig. 5E-F**). SCFAs commonly regulate gene expression through HDAC inhibition or GPCR activation^44^. However, keratinocytes express low levels of SCFA-sensing GPCRs^15^, and inhibition of downstream G protein signaling did not attenuate Bu-induced PDGFB expression (**Fig. S4C-E**), making GPCR-mediated signaling less likely. Although the HDAC inhibitor SAHA modestly induced PDGFB expression, the magnitude of induction was substantially lower than that observed with Bu, despite robust activation of control genes and induction of keratinocyte differentiation (**Fig. 5A-B, S4A-B**). These findings suggest that HDAC inhibition alone is insufficient to account for Bu-induced PDGFB expression. Bu also functions as a metabolic substrate and has been shown to promote oxidative phosphorylation and fatty acid oxidation in keratinocytes^16^. While PDGFB induction positively correlated with differentiation, differentiation itself appeared sensitive to both HDAC inhibition and histone acetylation, whereas PDGFB induction was preferentially dependent on acetyltransferase p300 activity (**Fig. 5A-B, 5G-I**). The uncoupling of differentiation and PDGFB expression observed in both the SAHA and calcium differentiation experiments further supports a model in which PDGFB induction relies primarily on HAT activity (**Fig. 5A-B, S3C**). Consistent with this model, Bu and propionate, but not acetate, induced PDGFB expression (**Fig. S3D-E**). This observation aligns with reports that Bu and propionate promote p300 acylation and activation, whereas acetate alone may primarily increase acetyl-CoA availability without directly activating p300^56^. Together, although our transcriptomic analyses indicated that Bu induced metabolic pathways associated with acetyl-CoA generation, further data suggest that p300 activation rather than acetyl-CoA accumulation alone is the key determinant of PDGFB induction.

Several other microbial metabolites have also been reported to promote collagen production. Indole-lactic acid (ILA) has been shown to restore dermal collagen in aged mice following fecal microbiota transplantation, and notably, topical application of an AhR inhibitor reversed this phenotype^37^. Although the cellular mechanism was not defined, this finding implies that epidermal signaling contributes to ILA-mediated collagen regulation. Consistent with this interpretation, we observed modest induction of TGFβ1 in keratinocytes treated with ILA (**Fig. 4A**), supporting a potential epidermal relay mechanism. Similarly, indole-acetic acid (IAA) has been reported to increase collagen expression in scleral fibroblasts through SP-1 activation^36^. We did not observe similar effects of IAA in human dermal fibroblasts (**Fig. 4A**), suggesting potential tissue-specific responses to microbial metabolites.

To further assess whether this epidermal relay mechanism extends beyond reductionist culture systems, we utilized explants derived from aged human skin to preserve native epidermal-dermal architecture (**Fig. 6A**). Consistent with our *in vitro* findings, topical Bu induced epidermal PDGFB expression, which could be attenuated by p300 inhibition (**Fig. 6B, D**). Increased epidermal PDGFB expression was accompanied by upward trends in dermal collagen transcriptional responses and supportive histological observations (**Fig. 6C-D**). The relatively modest collagen response compared with PDGFB induction may reflect differences in kinetics, with growth factor induction occurring more rapidly than extracellular matrix remodeling, particularly within the short 24-hour treatment window. Together, these findings further support a model in which microbial metabolite sensing by the epidermis regulates downstream dermal responses in intact human skin.

Targeting dermal fibroblasts to enhance collagen production is an attractive therapeutic strategy but is limited by the epidermal permeability barrier which prevents penetration of topical agents into the dermis^68^. Our findings suggest an alternative approach in which keratinocytes act as intermediaries that translate microbial metabolite exposure into dermal signaling. In this context, Bu-induced PDGFB expression in keratinocytes provides a mechanism by which epidermal stimulation can amplify downstream dermal collagen production (**Fig. 6E**). A similar epidermal-dermal PDGFB signaling axis has been reported previously, where reconstruction of the basement membrane increased keratinocyte-derived PDGF-BB and promoted dermal collagen deposition in an organotypic human skin model ^69^. In contrast, our findings identify microbial metabolites as upstream regulators of epidermal PDGFB expression, suggesting that microbiota-derived signals may also engage this pathway. Together, these findings support targeting keratinocytes as a feasible strategy to modulate dermal homeostasis.

Maintaining dermal collagen homeostasis is essential for skin integrity and function, and its disruption contributes to a range of pathological conditions, with additional implications for cosmetic and regenerative outcomes. Although platelet-rich plasma and PDGF injections are increasingly used in cosmetic and regenerative settings, their efficacy and safety remain variable and, in some contexts, controversial^70,71^. Leveraging microbiota and/or microbial metabolites to enhance dermal integrity may therefore represent a complementary and more physiologic approach, as microbial-derived signals are involved in endogenous skin regulation and barrier maintenance^3,72^.

Together, our findings support a model in which microbiota-derived Bu promotes epidermal-dermal crosstalk through PDGFB signaling to regulate dermal collagen homeostasis (**Fig. 6E**). This work expands the functional scope of the microbiota beyond immune and barrier regulation and highlights microbial metabolites as potential therapeutic modulators of dermal integrity.

## Materials and Methods

### Animal models

#### Husbandry conditions

All mouse procedures were conducted under protocols approved by the University of Pennsylvania Institutional Animal Care and Use Committee (protocols 804065 and 805700). Germ-free (GF) mice were bred and maintained in flexible film isolators in the Penn Gnotobiotic Mouse Facility (PGMF) at the University of Pennsylvania School of Veterinary Medicine. For experiments, GF mice were transferred to hermetically sealed isocages with individual ventilation and sterile food and water to maintain GF status and housed in a BSL II facility within the PGMF for the duration of the study. Specific pathogen-free, conventionally raised (CR) C57BL/6J mice were purchased from The Jackson Laboratory (strain 000664) and housed in a BSL I facility within the PGMF. Upon arrival, CR mice were acclimated for one week prior to experimentation. Unless otherwise noted, all adult mice used in this study were 12 weeks old at the experimental endpoint and were age- and sex-matched within each experiment.

#### Colonization of germ-free mice

Samples for murine dermal RNA sequencing were derived from mice generated as part of a previously published study (Figure 4 of Uberoi et al., *Cell Chemical Biology*, 2025^6^). For colonization, bacteria comprising the skin microbial consortium Flowers’ Flora 50 (FF) were individually cultured and combined at 10^7 CFU per microbe. The FF mixture was inoculated onto cage bedding every other day for four weeks. Tail dermis was harvested at the experimental endpoint for downstream analyses.

#### Timed mating and antibiotic treatment

Timed matings were established by pairing adult females with males overnight. Embryonic day 0.5 (E0.5) was defined by detection of a copulation plug using a sterile vaginal probe the following morning. Pregnant dams received ceftriaxone (250 mg/mL in sterile deionized water; 0.2 mL per dose; GoldBio C-781-1) or sterile deionized water as vehicle control by oral gavage from E7.5 to E9.5 and again from E11 to E16, with a rest period between dosing windows. Embryos were harvested at E17 for downstream analyses.

### Cell and tissue culture

#### Primary human cells

Primary cultures of human keratinocytes and fibroblasts were obtained from neonatal foreskins through the Skin Translational Research (STaR) Core of the Penn Skin Biology and Diseases Resource-based Center (SBDRC). All experiments were conducted with keratinocytes at passage 2 and fibroblasts at less than 4. Keratinocytes were cultured in a keratinocyte growth media: 50% Media 154 (Life Technologies M154500), 50% Keratinocyte SFM with its supplements (Life Technologies, 17005042), 1% HKGS supplement (Life Technologies, S0015). Fibroblasts were culture in high glucose DMEM (Gibco 11965118) with 10% FBS, 1% sodium pyruvate (Gibco 11360-070) and 1% antibiotic/antimycotic (Invitrogen 15240062). Cells were split when they were less than 80% confluent. Cells were washed with 1X PBS and trypsinized with 0.25% Trypsin-EDTA (Gibco 25200-056) for 5 minutes, trypsin was inactivated using trypsin inhibitor (Sigma-Aldrich T6414) and cell suspension was centrifuged. Cell pellet was suspended in culture media and seeded as per experimental design. Cells were maintained at 37 °C in an atmosphere of 5% CO2 with humidity.

#### Monolayer cell treatments

Primary keratinocytes and fibroblasts were seeded and allowed to adhere overnight prior to treatment. Cells were treated with metabolites or inhibitors at the indicated concentrations and durations for each experiment. To induce keratinocyte differentiation, growth media was supplemented with calcium to a final concentration of 1.2 mM one day prior to butyrate treatment. Cells were subsequently treated with butyrate for 24 hours before harvest. For conditioned media experiments, primary fibroblasts were treated with HEE-derived conditioned media at a final concentration of 20% in fibroblast growth media for 24 hours before harvest.

#### Human epidermal equivalents

Human epidermal equivalents (HEEs) were generated following a workflow adapted from Pavel et al., *JID Innovations*, 2021^73^. Primary keratinocytes were seeded onto transwell inserts (Millipore PTHT12H48) and allowed to proliferate in keratinocyte growth media for three days before switching to differentiation media consisting of 70% Epithelial 3D Airlift Media (CnT-PR-3D) and 30% high-glucose DMEM. Cultures were air-lifted the following day and maintained at the air-liquid interface for five days, with differentiation media supplied to the basal chamber.

For metabolite screening and suberoylanilide hydroxamic acid experiments, compounds were added at the time of air lift and refreshed every two days with differentiation media. For G protein inhibitor experiments, inhibitors were added one day prior to treatment with 1 mM butyrate, and cultures were harvested after 24 hours. For other inhibitor experiments, inhibitors and butyrate were added one day prior to harvest.

#### Human skin explants

Human skin explants were generated from aged human skin (>65 years old) obtained from Mohs surgery through the STaR Core of the Penn SBDRC. Subcutaneous fat was removed and 6 mm punch biopsies were generated. Explants were cultured on transwell inserts (Corning CLS3401) in equilibration media consisting of a 1:6 mixture of heat-inactivated FBS and high-glucose DMEM supplemented with 1% antibiotic-antimycotic overnight. The following day, 5 µL of 2 mM butyrate with or without 5 µM A485 (MedChemExpress HY-107455) formulated in a 2:8 mixture of propylene glycol and isopropanol was topically applied to the explants. Explants were subsequently cultured for 24 hours in treatment medium consisting of 50% Epithelial 3D Airlift Media and 50% high-glucose DMEM. Following treatment, explants were bisected. One half was immediately fixed for histologic analysis, while the remaining half was incubated in dispase solution (1 U/mL in PBS, Roche 37045800) overnight at 4 °C for epidermal-dermal separation prior to transcriptomic analysis.

#### Reagents and treatments

Sodium butyrate (13121), indole-3-acetic acid (16954), indole-3-lactic acid (37534), indole-3-propionic acid (28821), YM-254890 (29735), Doramapimod (10460), AX-15836 (28774) and PD0325901 (13034) were purchased from Cayman. Ascorbic acid (A4544) and sodium propionate (P1880) were purchased from Sigma-Aldrich. Pertussis Toxin was purchased from Hello Bio (HB4729). JNJ-10198409 (6976) and suberoylanilide hydroxamic acid (4652) was from Tocris.

#### Primary murine cells

Primary murine dermal fibroblasts were isolated from ear skin. Ear tissue was split into dorsal and ventral sheets by manually separating the skin from the underlying cartilage. Tissue was incubated in dispase solution (2 U/mL in PBS) overnight at 4 °C. The dermis was separated from the epidermis the following day and incubated in 3.5 mg/mL collagenase I for 1 hour at 37 °C with shaking. Following digestion, fibroblast growth media as described above was added to the mixture. Liberated cells were filtered through a 100 µm cell strainer, pelleted by centrifugation, and plated in fibroblast growth media as described above.

### Dermal RNA sequencing

#### Tissue collection and processing

Tail skin from CR, GF, and FF-colonized mice was dissected from the tail bone and incubated in dispase solution overnight at 4 °C. The dermis was separated from the epidermis the following day and immediately snap frozen and stored at −80 °C until processing.

Frozen dermal tissue was cryo-pulverized using a CP02 cryoPrep Automated Dry Pulverizer (Covaris), transferred to Lysing Matrix D tubes (MP Biomedicals), and homogenized in 1 mL TRIzol (Invitrogen 15596026) using a Mini-Beadbeater (BioSpec). RNA was purified by phenol-chloroform extraction followed by on-column DNase digestion (Qiagen 79254).

Library preparation and sequencing were performed by the Children’s Hospital of Philadelphia High Throughput Sequencing Core. Libraries were constructed using Illumina Stranded Total RNA Prep with Ribo-Zero Plus and sequenced on a NextSeq 1000 using P2 flow cells to a depth of approximately 20–25 million paired-end reads per sample (50 bp).

#### Analysis

RNA-seq data were analyzed using a workflow adapted from the DIYtranscriptomics framework^74^. Reads were pseudoaligned to Genome Reference Consortium Mouse Build 39 (GRCm39) using kallisto (version 0.51.0)^75^. Transcript abundances were imported into R (version 4.5.1) and summarized to the gene level using tximport (version 1.36.1) with length-scaled transcripts per million and gene annotations from EnsDb.Mmusculus.v79^76^. Gene-level counts were further processed using edgeR (version 4.6.3) to calculate counts per million, filter lowly expressed genes, and normalize expression across samples^77^. Normalized expression values were analyzed using limma (version 3.64.3) following voom transformation, with linear modeling and empirical Bayes moderation used to identify differentially expressed genes^78^. Pairwise differential expression between experimental conditions was assessed using specified contrasts. Differentially expressed genes (DEGs) from CR versus GF comparisons were further annotated using the murine Matrisome database (revision 2014) from the Matrisome Project^24^.

### Reanalysis of published epidermal RNA sequencing data

Raw sequencing data from the previously published epidermal RNA-seq dataset^23^ (GSE162925) were obtained and reprocessed using the RNA-seq workflow described above. Reads were pseudoaligned to the mouse reference genome (GRCm39) and analyzed using kallisto, tximport, edgeR, and limma as described for dermal RNA-seq. P values were adjusted for multiple testing using the Benjamini-Hochberg method, and genes with an adjusted P value below 0.05 were considered differentially expressed. Dermal and epidermal differentially expressed genes were integrated and annotated using the ligand-receptor database from OmniPath^43^.

### Human epidermal equivalent RNA sequencing

Human epidermal equivalents with short term (1 day) and long term (5 day) butyrate treatment were harvested by excising the transwell inserts with attached membranes and homogenized in RNeasy Mini Kit lysis buffer (Qiagen 74106) by bead beating. RNA was extracted using the RNeasy Mini Kit with on-column DNase digestion (Qiagen 79254) according to the manufacturer’s instructions. Library preparation and sequencing were performed as described for dermal RNA-seq. Data were processed and analyzed using the same computational workflow described above. Reads were pseudoaligned to Human GRCh38. P values were adjusted using the Benjamini-Hochberg method and genes with adjusted P value below 0.05 were considered differentially expressed.

### RNA extraction and cDNA synthesis for qPCR

Fibrous tissues were cryo-pulverized prior to homogenization as described above. Non-fibrous tissues were homogenized directly by bead beating. Total RNA from murine tissues was purified by phenol-chloroform extraction followed by in-solution DNase digestion (Promega M6101) according to the manufacturer’s protocol. RNA from cultured cells and human explants was extracted using the RNeasy Mini Kit (Qiagen 74106) with on-column DNase digestion (Qiagen 79254).

Complementary DNA was synthesized using SuperScript III Reverse Transcriptase (Invitrogen 18080044). Quantitative PCR was performed using TaqMan Fast Advanced Master Mix (Applied Biosystems 4444557) and TaqMan Gene Expression Assays for the following targets: *Col1a1* (Mm00801666_g1), *Loxl3* (Mm01184865_m1), *Gulo* (Mm00626646_m1), *Pdgfb* (Mm00440677_m1), *Rplp2* (Mm03059047_gH), *Tgfb1* (Mm00441724_m1), *COL1A1* (Hs00164004_m1), *LOXL3* (Hs01046941_g1), *PDGFB* (Hs00966522_m1), *IVL* (Hs00846307_s1), *AQP3* (Hs00185020_m1), *KRT14* (Hs00265033_m1), *KRT10* (Hs00166289_m1), *LOR* (Hs01894962_s1), *FLG* (Hs00856927_g1), and *RPLP0* (Hs04189669_g1). Reactions were run on a QuantStudio 7 Pro Real-Time PCR System (Thermo Fisher Scientific).

### Picrosirius red staining, visualization and analysis

#### Staining and visualization

Adult murine dorsal skin and ear tissue, as well as embryonic bisected trunks, were collected and fixed overnight in 4% paraformaldehyde, then transferred to 70% ethanol prior to submission to the Cutaneous Phenomics and Transcriptomics (CPAT) Core of the Penn SBDRC. Tissues were paraffin-embedded, sectioned at 5 µm, and stained with picrosirius red (PSR; Polysciences 24901) by the CPAT Core according to the manufacturer’s protocol.

Bright field and polarized light images were acquired using a Leica DM6 B upright microscope. For adult tissues, entire sections were tile-scanned using a 10X objective and stitched using LAS X software (version 3.7.6.25997). For embryonic tissues, two representative fields per section were imaged using a 20X objective.

#### Analysis

Images were rotated and cropped in Adobe Photoshop (version 24.7.0) and quantified in ImageJ (version 2.14.0/1.54f). For adult tissues, polarized light images were separated into red, green, and blue channels, and 8-bit red channel images were used for downstream analyses. Embryonic tissues were converted to 8-bit grayscale without channel separation due to mixed collagen birefringence. Dermal regions of interest were manually outlined based on PSR signal boundaries, the epidermal-dermal junction, hair follicles, and the panniculus carnosus.

Dermal thickness was quantified by averaging five measurements per section for each mouse. Collagen staining intensity was quantified from histogram-derived pixel intensity values in ImageJ and analyzed in RStudio (version 2025.09.1+401). For adult tissues, intensity thresholds were defined based on the distribution of pixel values across samples, with a high-intensity cluster observed in the 200–255 range. Thus, pixels with intensity values greater than 200 were classified as strong signals and normalized to the total pixel number within each dermal region. For embryonic tissues, where a distinct high-intensity cluster was not observed, pixels with intensity values greater than 50 were included for intensity calculations.

For collagen fiber metrics, processed 8-bit red channel images of dermal regions were binned to reduce image size and analyzed using the TWOMBLI workflow according to the user tutorial^28^.

### Histology and immunofluorescence

HEEs and human skin explants were fixed and processed by the Penn SBDRC CPAT Core as described above. Sections were subjected to hematoxylin and eosin (HCE) staining or immunofluorescence staining. HCE-stained sections were imaged using a Keyence BZ-X710 microscope. For immunofluorescence staining of PDGFB (ABclonal A1195) and procollagen type I (RCD Systems AF6220) nuclei were counterstained with DAPI, and images were acquired using the Leica microscope.

### Hydroxyproline assay

Murine skin and bone tissues were cryo-pulverized prior to homogenization as described above. Intestinal tissues and cell culture samples were homogenized directly by bead beating. All samples were processed in ultrapure water prior to hydrolysis. Skin, intestinal, and cell culture samples were hydrolyzed in 5 M sodium hydroxide overnight at 95 °C. Bone samples were hydrolyzed in 5 M hydrochloric acid overnight at 95 °C to allow simultaneous decalcification. Following neutralization the next day, hydroxyproline assays were performed according to the manufacturer’s protocol (Sigma-Aldrich MAK463). Total protein equivalents of neutralized hydrolysates were estimated using a bicinchoninic acid (BCA) assay (Thermo-Scientific 23227) with hydrolyzed bovine serum albumin standards processed through the same hydrolysis procedure. Hydroxyproline values were normalized to the corresponding BCA signal for each sample.

### Epidermal tissue ELISA

Adult murine dorsal skin was collected and incubated in dispase solution overnight at 4 °C. The epidermis was separated from the dermis the following day and homogenized by bead beating in RIPA buffer (Research Products International R26200) supplemented with protease and phosphatase inhibitor cocktails (ThermoFisher 78442). Latent TGFβ (BioLegend 594509) and PDGFB (RCD Systems MBB00) concentrations were measured using ELISA kits according to the manufacturers’ instructions. Total protein concentration was determined using a BCA assay. ELISA values were normalized to the corresponding total protein content for each sample.

### Data availability

The RNA sequencing datasets generated in this study, including dermal and HEE transcriptomic datasets, have been deposited in the NCBI Gene Expression Omnibus under accession number GSE334945 and will be publicly available upon publication.

