## Supplemental Methods and Figure S1-S4 for "Microbiota-derived butyrate promotes dermal collagen production through epidermal-dermal crosstalk"

**Title**


**Supplementary Contents**

- Supplementary Materials and Methods
- Supplementary Reference
- Supplementary Figures and Figure Legends

### Supplementary Materials and Methods

#### Scratch closure assay

Primary murine dermal fibroblasts derived from CR and GF mice were seeded using two-well culture inserts (Ibidi 80209) placed in 12-well plates at 3 × 10^5 cells/mL to generate standardized cell-free gaps, following the manufacturer's instructions. Cells were allowed to adhere overnight, after which inserts were removed, and cells were maintained in fibroblast growth media as described above. Wound gaps were imaged at 0, 12, and 24 hours using the Keyence microscope. Images were acquired at the same locations for each well at all time points. Gap area was quantified using the Wound Healing Size Tool plugin in ImageJ following the developer's tutorial[1]. For each well, gap size at each time point was normalized to the corresponding baseline measurement at 0 hours. Linear regression was used to fit the rate of gap closure, and differences between groups were assessed using analysis of covariance (ANCOVA).

#### Western blot analysis

Cells were lysed in RIPA buffer supplemented with protease and phosphatase inhibitors. Protein concentrations were determined using a BCA assay. Equal amounts of protein were separated by SDS-PAGE and transferred to PVDF membranes (Bio-Rad). Membranes were blocked using Intercept (TBS) Blocking Buffer (LICORbio 927-60001) and incubated with anti-COL1A1 antibodies (Invitrogen PA5-29569) overnight at 4 °C. After washing, membranes were incubated with IRDye secondary antibodies (LICORbio 926-68073) and imaged using an Odyssey imaging system. GAPDH (Cell Signalling 2118) was used as a loading control. Band intensities were quantified using ImageJ.

#### In-solution picrosirius red assay

Soluble collagen from fibroblast cultures was quantified using an in-solution PSR assay[2]. Primary murine dermal fibroblasts derived from CR and GF mice were cultured in fibroblast growth media, and conditioned media were collected after 5 days. Conditioned media were incubated with PSR solution (Polysciences 24901) to allow collagen binding, followed by centrifugation to pellet collagen-bound dye. Pellets were washed to remove unbound dye and subsequently dissolved in alkaline solution. Absorbance was measured using a microplate reader, and collagen levels were quantified based on optical density 570 nm measurements.

#### Matrix swapping scratch assay

Cell-derived extracellular matrix (ECM) preparation was adapted from Hellewell et al.[3]. Donor fibroblasts were cultured for 5 days prior to decellularization with 50 mM ammonium hydroxide (Sigma-Aldrich 09859) for 20 minutes at room temperature, followed by PBS washes to ensure complete removal of donor cells. Recipient fibroblasts were then seeded onto donor-derived ECM using two-well culture inserts as described above. Following insert removal, recipient fibroblasts were maintained in fibroblast growth medium supplemented with 20% donor-derived conditioned media to reconstitute soluble components of the donor matrisome. Scratch assays were subsequently performed as described above.

### Supplementary Figures and Figure Legends

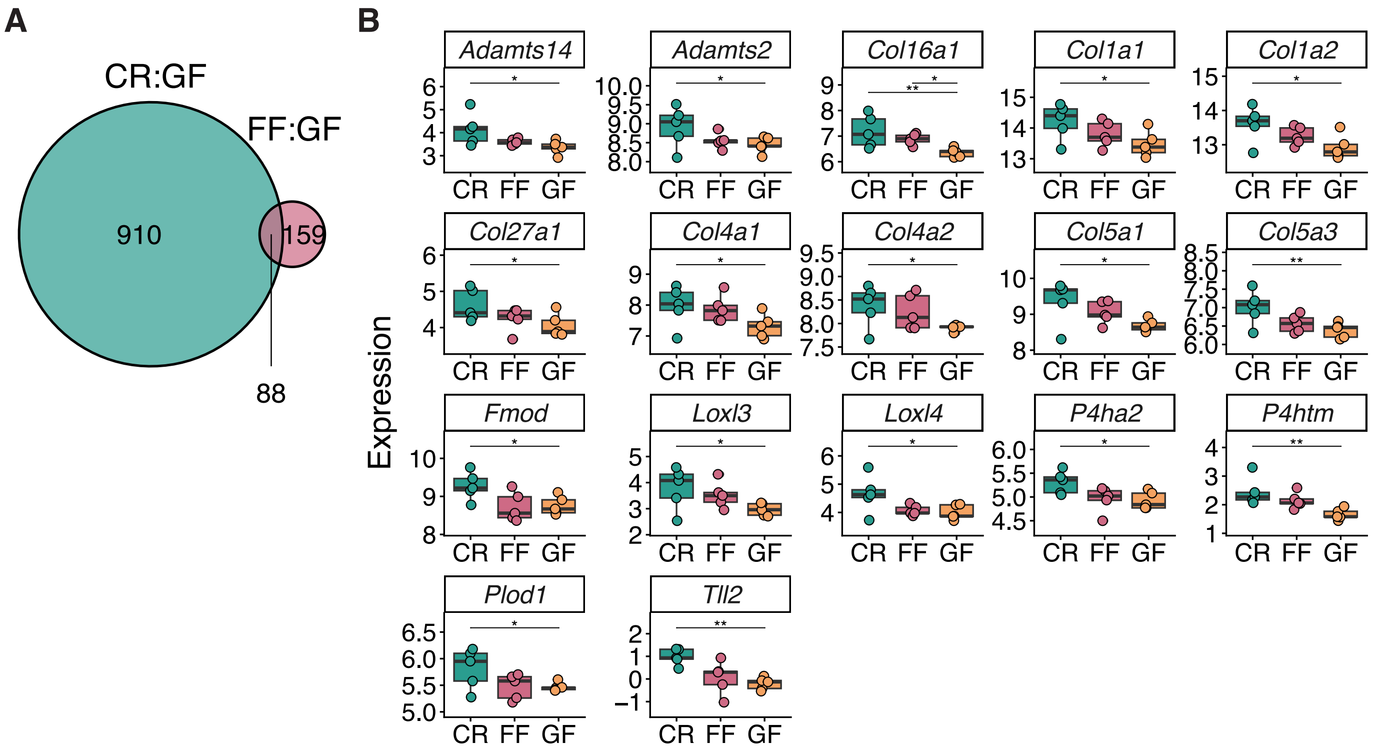


**Figure S1. Short term skin colonization with human skin commensals (FF) partially restores dermal matrisome gene expression**

(A) Venn diagram showing the number of overlapping DEGs identified in comparisons of CR versus GF and FF versus GF dermis.

(B) Expression levels of collagen biosynthesis genes enriched in CR dermis across CR, FF, and GF groups. Data were extracted from dermal RNA-seq.

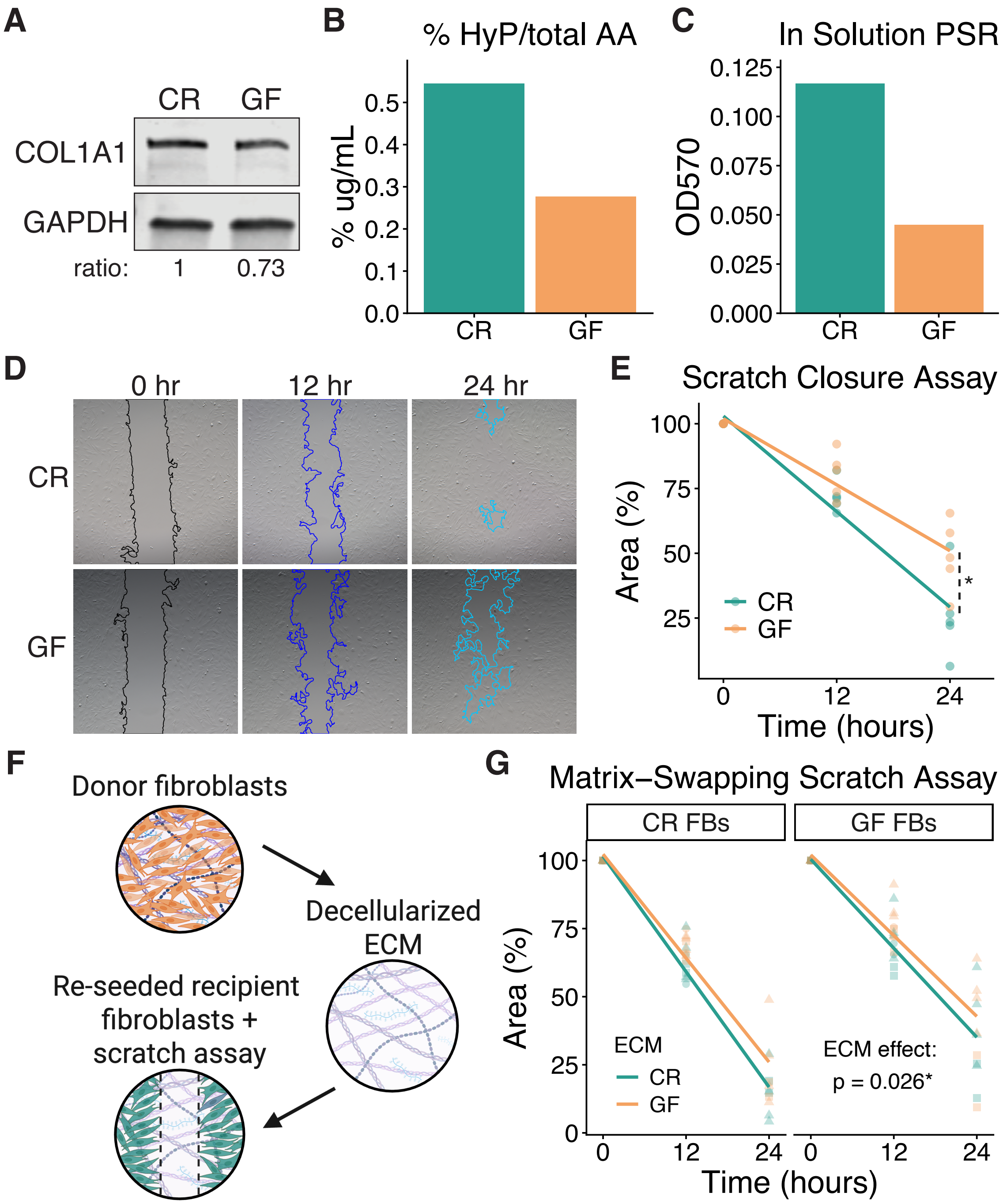


**Figure S2. Reduced collagen expression and diminished migratory capacity of germ-free fibroblasts**

(A-C) Collagen production was assessed using pooled dermal fibroblasts isolated from ear skin of CR and GF mice. N = 5 mice per group.

(A) Western blot showing COL1A1 and GAPDH expression in fibroblast cultures. COL1A1 levels were quantified relative to GAPDH.

(B) HyP content of fibroblast cultures normalized to total AA measured by BCA assay.

(C) In solution PSR staining of conditioned media from fibroblast cultures.

(D-E) Fibroblast migration was assessed using fibroblasts derived from individual CR and GF mice. N = 5 mice per group.

(D) Representative images of fibroblast migration assays across 0, 12, and 24 hours after scratch creation.

(E) Gap area relative to initial area over time, fit using a linear model. Each dot represents one mouse. *, $P < 0.05$ by ANCOVA.

(F-G) Matrix-swapping scratch assays were performed using pooled dermal fibroblasts isolated from ear skin of CR and GF mice. N = 5 mice per group.

(F) Workflow illustrating matrix swapping combined with scratch assay. Recipient fibroblasts were seeded onto ECM scaffolds generated from decellularized 5-day donor fibroblast cultures prior to scratch migration assays.

(G) Gap area relative to initial area over time, fit using a linear model (area ~ cells * ECM), faceted by recipient fibroblasts and colored by ECM donor fibroblasts. P-value determined by ANCOVA. Each dot represents one technical replicate, and dot shapes denote independent experimental repeats.

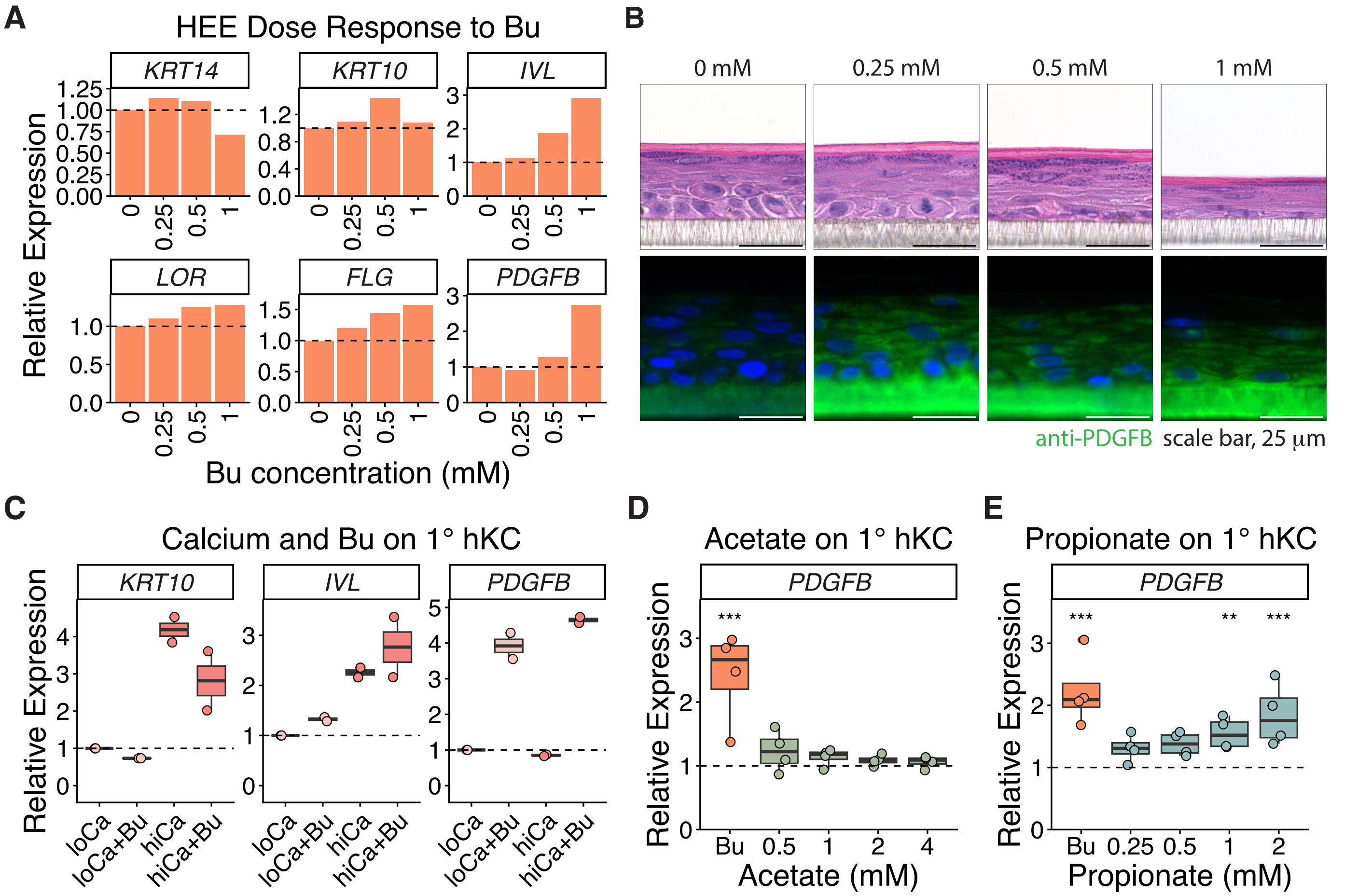


**Figure S3. Dose-dependent induction of epidermal PDGFB by butyrate is independent of keratinocyte differentiation**

(A) *KRT14*, *KRT10*, *IVL*, *LOR*, *FLG*, and *PDGFB* mRNA levels normalized to *RPLP0* from HEEs treated with vehicle or indicated concentrations of Bu. Dashed lines denote vehicle control levels normalized to 1.

(B) Hematoxylin and eosin stained (top) and PDGFB immunostained (bottom) HEEs treated as indicated. Nuclei were counterstained with DAPI. Scale bar, 25 µm.

(C) *KRT10*, *IVL*, and *PDGFB* mRNA levels normalized to *RPLP0* from 1° hKC primed for 24 hours in either low calcium (loCa, no added calcium) or high calcium (hiCa, 1.2 mM) media and subsequently treated with vehicle or 1 mM Bu for 24 hours under the same calcium conditions. Dashed lines denote loCa vehicle control levels normalized to 1 for each donor. N = 2 donors.

(D-E) *PDGFB* mRNA levels normalized to *RPLP0* from 1° hKC treated for 24 hours with vehicle, 1 mM Bu, or indicated concentrations of acetate (D) or propionate (E). Dashed lines denote vehicle control levels normalized to 1 for each donor. N = 4 donors. **, $P < 0.01$; ***, $P < 0.001$ by linear mixed effects model with donor as a random effect.

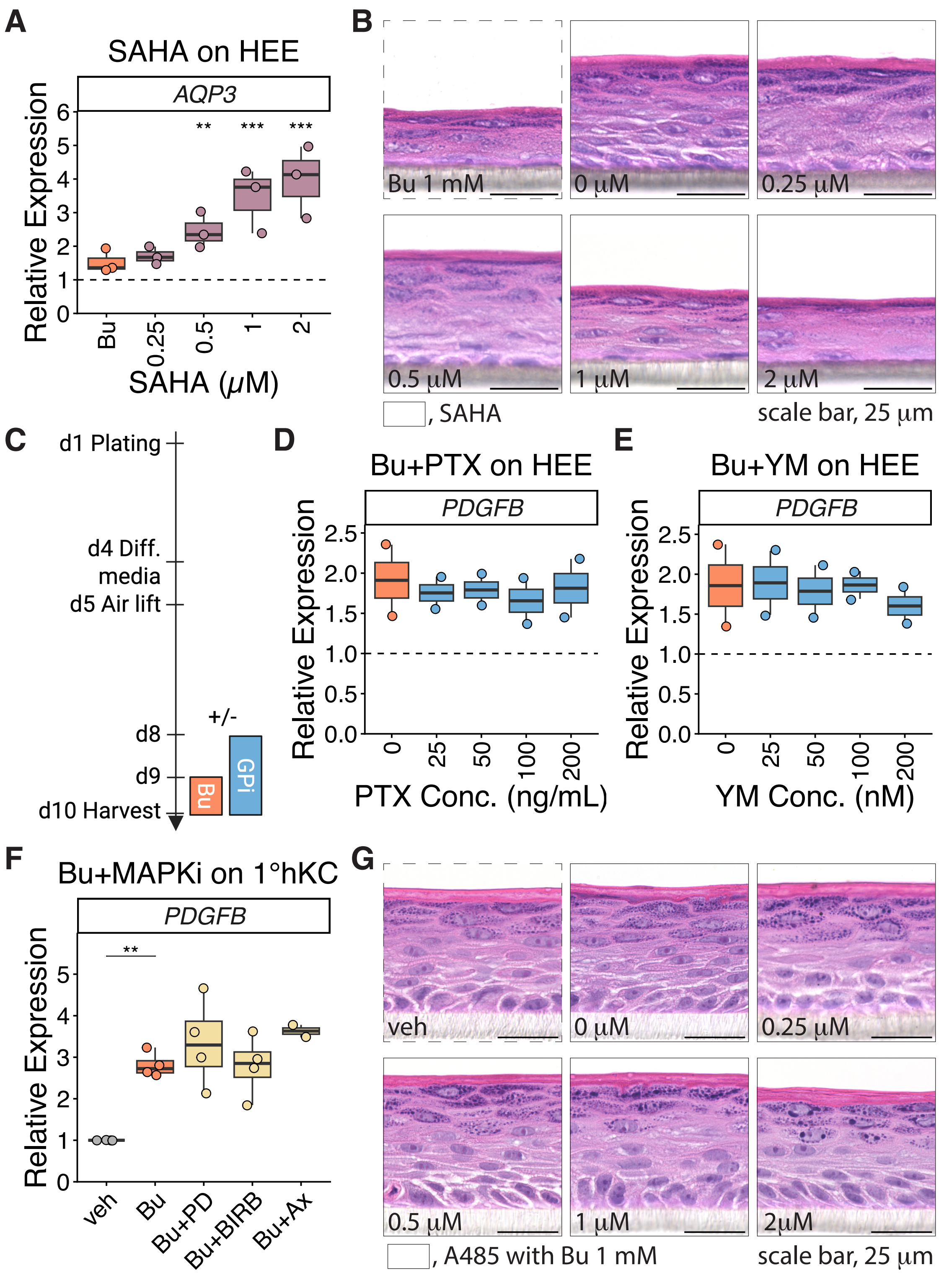


**Figure S4. Butyrate-induced PDGFB expression is not mediated by HDAC inhibition or GPCR/MAPK signaling**

(A) *AQP3* mRNA levels normalized to *RPLP0* from HEEs treated with vehicle, Bu, or indicated concentrations of SAHA. The dashed line denotes vehicle control levels normalized to 1 for each donor. N = 3 donors.

(B) H&E stained HEEs treated as indicated. Dashed boxed region denotes HEE treated with Bu; solid boxed regions denote HEEs treated with vehicle or SAHA. Scale bar, 25 µm.

(C) Workflow illustrating pretreatment of HEEs with G protein inhibitors (GPi) one day prior to the addition of 1 mM Bu.

(D-E) *PDGFB* mRNA levels normalized to *RPLP0* from HEEs pretreated with (D) a Gi/o inhibitor (pertussis toxin, PTX) or (E) a Gq/11 inhibitor (YM-254890, YM) at indicated concentrations and subsequently treated with 1 mM Bu under the same PTX/YM concentrations. Dashed lines denote vehicle control without Bu or PTX/YM normalized to 1 for each donor. N = 2 donors.

(F) *PDGFB* mRNA levels normalized to *RPLP0* from 1° hKC treated for 24 hours with vehicle, 1 mM Bu, Bu plus 200 nM MEK1/2 inhibitor (PD0325901, Bu+PD), Bu plus 1 μM p38 inhibitor (Doramapimod/BIRB, Bu+BIRB), or Bu plus 1 μM ERK5 inhibitor (AX-15836, Bu+AX). Data are expressed relative to the corresponding vehicle control for each donor. N = 4 donors for PD and BIRB; N = 2 donors for AX. **, $P < 0.01$ by linear mixed effects model with donor as a random effect.

(G) H&E stained HEEs treated with 1 mM Bu and indicated concentrations of A485. Dashed boxed region denotes vehicle (veh) treated HEE; solid boxed regions denote HEEs treated with 1 mM Bu and increasing concentrations of A485. Scale bar, 25 μm.
